# The processing of color preference in the brain

**DOI:** 10.1101/361006

**Authors:** Chris Racey, Anna Franklin, Chris M. Bird

## Abstract

Decades of research has established that humans have preferences for some colors (e.g., blue) and a dislike of others (e.g., dark chartreuse), with preference varying systematically with variation in hue (e.g., Hurlbert & Owen, 2015). Here, we used functional MRI to investigate why humans have likes and dislikes for simple patches of color, and to understand the neural basis of preference, aesthetics and value judgements more generally. We looked for correlations of a behavioural measure of color preference with the blood oxygen level-dependent (BOLD) response when participants performed an irrelevant orientation judgement task on colored squares. A whole brain analysis found a significant correlation between BOLD activity and color preference in the posterior midline cortex (PMC), centred on the precuneus but extending into the adjacent posterior cingulate and cuneus. These results demonstrate that brain activity is modulated by color preference, even when such preferences are irrelevant to the ongoing task the participants are engaged. They also suggest that color preferences automatically influence our processing of the visual world. Interestingly, the effect in the PMC overlaps with regions identified in neuroimaging studies of preference and value judgements of other types of stimuli. Therefore, our findings extends this literature to show that the PMC is related to automatic encoding of subjective value even for basic visual features such as color.

## Introduction

Over the last century, scientists have provided converging evidence that humans have reliable and systematic preferences for some colors over others (see Hurlbert & Owen, 2015; Palmer, Schloss & Sammarinto, 2013, for reviews). Whilst individuals vary in their color preferences, colors such as blue are commonly the most liked and other colors such as yellowy-green the most disliked. In addition, when preferences are plotted against variation in hue, this reveals a systematic hue preference curve which rises steadily as hues get bluer and less yellow, although there is also some interaction with lightness and saturation (e.g., saturated yellow is more commonly liked than dark yellow: Palmer & Schloss, 2010). This pattern of color preference has been relatively stable over time. For example, such preferences were revealed in the earliest studies of color preference dating as far back as the 19^th^ century (Jastrow, 1893 cited in Ling, Hurlbert & Robinson, 2006). In addition, although there is some cultural variation in the hue preference curve (e.g., Taylor, Clifford & Franklin, 2013), the general pattern appears to be consistent across industrialised cultures - some have even claimed that some aspects of color preference are ‘universal’ (Hurlbert & Ling, 2007).

The presence of reliable and systematic color preferences raises the question of why some colors are liked by humans more than others. What is it about color that ‘holds’ affect? (Zajonc, 1980). Potential answers to this question have been provided by behavioural studies that have identified relationships between color preference and various measures such as the ease of naming different colors (e.g., Àlvaro, Moreira, Lillo & Franklin, 2015), the valence and number of color-object associations (Palmer & Schloss 2010; Schloss & Palmer, 2017; Taylor & Franklin, 2012), and the sensory mechanisms of early color encoding (Hurlbert & Ling, 2007). Here we propose that identifying regions of the brain where BOLD activity relates to color preferences could provide leverage in understanding color preference, as well as furthering understanding of the neural basis of human preference, aesthetics and abstract higher level judgements more generally.

There have been a few previous fMRI studies of different aspects of color aesthetics (Ikeda, Matsuyoshi, Sawamoto, Fukuyama, & Osaka, 2015; Johnson, Lowery, Kohler, & Turetsky, 2005; Kim, Song, & Jeong, 2012). However, none of these enable us to reliably identify the regions of the brain that are more active when people view colors that they like than those they dislike. For example, the orbitofrontal cortex and amygdala are associated with judgments of color harmony, yet harmony is a process distinct from color preference (Ikeda et al., 2015). Anterior medial prefrontal cortex (AMPFC) and retrosplenial cortex (RSC) regions are more active when people make preference judgments compared to similarity judgements about color (Johnson et al., 2005), but it does not necessarily follow that these regions are associated with how much colors are liked. Only one prior study has aimed to identify brain regions where activation on viewing colors is associated with preference for those colors (Kim et al., 2012). However, this study only contrasted activity while viewing a liked color (green) with black: a comparison that is likely to be driven mainly by the difference in lightness. Moreover, the effects are reported at lenient thresholds well below those standardly used in the field. It is therefore impossible to draw firm conclusions from this study.

The current fMRI study aims to identify brain regions associated with color preference: what regions of the brain have stronger activation on viewing a color the more a color is liked? We look for correlations of a behavioural measure of color preference with brain activity when people passively view those colors. Based on two distinct areas of research concerned with color preference (e.g., Àlvaro et al., 2015; Hurlbert & Ling, 2007; Palmer & Schloss, 2010) and the neural basis of value or aesthetic judgments (e.g., Grueschow, Polania, Hare, & Ruff, 2015; Vessel, Sterr & Rubin, 2013), we make two sets of predictions.

The first set of predictions is based on the identified relationships between color preference and color naming (Àlvaro et al., 2015), color-object associations (e.g., Palmer & Schloss, 2010) and the sensory mechanisms of color vision (Hurlbert & Ling, 2007). We localise regions of the brain that potentially underpin these relationships, using localisers for color naming, object perception (Grill-Spector, Kourtzi, & Kanwisher, 2001), and color selective regions of visual cortex (Lafer-Sousa & Conway, 2013). The prior behavioural evidence predicts a relationship of the blood oxygen level-dependent (BOLD) response in these regions when viewing colors with how much those colors are liked.

We make a second set of predictions based on prior studies of the neural basis of value and aesthetic judgements (a research area that some have termed ‘neuroaesthetics’, (Chatterjee, 2011; Conway & Rehding, 2013). Prior neuroimaging studies of judgements of the beauty of stimuli such as faces (Chatterjee, Thomas, Smith, & Aguirre, 2009), music (Tomohiro Ishizu & Zeki, 2011) or art (e.g., Vessel et al., 2013), have implicated a number of regions, particularly those included in the brain’s so-called default mode network (DMN; Raichle et al., 2001). Of these, the orbitofrontal cortex in particular, has been identified in several studies and suggested to be the ‘beauty centre’ of the brain (Ishizu & Zeki, 2013; Ishizu & Zeki, 2011). However, there has been debate about the extent to which this region is activated by the actual act of making a judgement of beauty or value rather than being related to these implicitly or automatically (Conway & Rehding, 2013; Tomohiro Ishizu & Zeki, 2014). A related literature on subjective value judgments (how much is this stimulus valued by you?) has suggested that another hub of the DMN, the posterior midline cortex (including the posterior cingulate cortex and precuneus), underlies automatic subjective value when value judgments about the stimuli are not explicitly being made (Grueschow et al., 2015).

In the current study, BOLD activity is measured when participants passively view colors and are engaged in a task unrelated to the colors. We correlate activity with a behavioural measure of color preference taken at the end of the experiment. In this way, we aim to identify the regions of the brain related to implicit and automatic color preferences rather than the actual process of judging preference. If we identify regions also implicated in judgements of value or beauty in prior work (e.g., Chatterjee, 2011), this would provide evidence that these regions are also related to implicit judgments, and that the involvement of these regions extends to color. Effects in posterior midline cortex would suggest that these regions underpin the computation of automatic subjective value across stimulus domains and different types of value.

## Method

### Subjects

Twenty-one healthy right-handed volunteers (11 males; 10 females; mean age: 26±4.4 years) participated. All were native English speakers, had normal or corrected-to-normal vision, and were screened for color vision deficiencies (Red-Green, Ishihara, 1983; Tritan, City 2nd Ed., Fletcher, 1980). Each participant gave written informed consent and the study was approved by the Research Governance and Ethics committee of the Brighton and Sussex Medical School and the European Research Council Executive Agency, conforming to the Code of Ethics of the World Medical Association (Declaration of Helsinki). Participants received a financial compensation of £20 for taking part in the study. All data were anonymized.

### Stimuli

There were 24 chromatic stimuli that were close approximations of the saturated (S), light (L), and dark (D) versions of 8 hues from the Berkeley Color Project (Palmer and Schloss, 2010), with minor adjustment to fit within the gamut of the stimulus display. The saturated set comprised of maximally saturated good examples of red (R), orange (O), yellow (Y), chartreuse (H), green (G), cyan (C), blue (B), and purple (P). The light and dark set stimuli had the same hue angles as the saturated set, but chroma (distance from the gray background in CIELUV color space) was set to approximately half of that for the corresponding hues in the saturated set. The dark set was 30% darker, and the light set was 20% lighter than the saturated stimuli (in CIE L*). There were also five achromatic colors (d65 grey) at evenly spaced steps of lightness (in CIE L*) from the minimum (black) to maximum (white) lightness renderable. See figure 1 for representations of the stimuli and their position in a perceptual color space, and table S1 in the supplementary section for the luminance and chromaticity co-ordinates of the stimuli. All stimuli were presented as square patches in the central 2° of the visual field on a grey background based on the standard d65 illuminant (x=0.3137, y=0.3214, 225.8 cd/m^2^). Visual stimuli were projected from a calibrated 3LCD video projector (Sony VPL-FE40) onto an screen outside of the scanner, which participants could see via a coil-mounted mirror. The chromaticity coordinates for the screen rendered stimuli were verified with a Spectrascan PR-655 spectroradiometer measuring from outside of the MRI bore via a system of mirrors.

**Figure 1.**
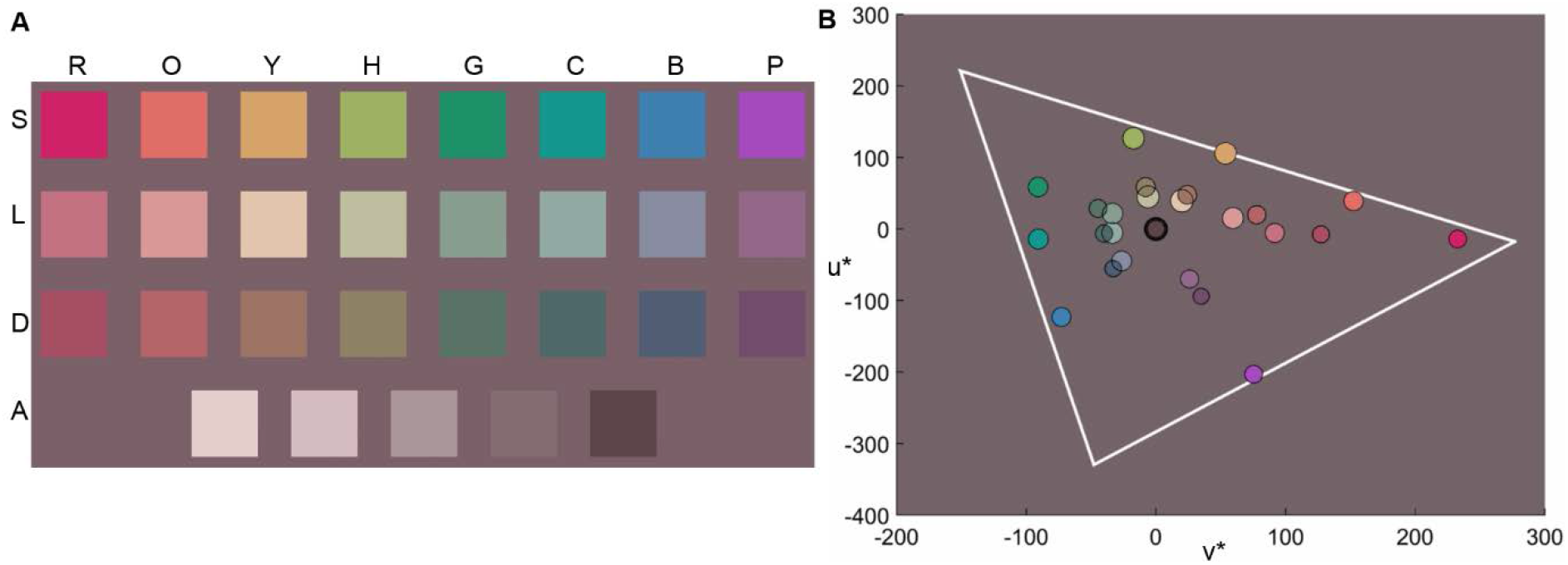
Examples of the stimuli and their position in color space. (**A**) Approximate rendering of saturated (S), light (L), dark (D), & achromatic (A) versions of red (R), orange (O), yellow (Y), chartreuse (H), green (G), cyan (C), blue (B) and purple (P) hues presented on the background grey used in the experiment. Colors are shown here for illustrative purposes and are only an approximation of those in the experiment because of variation in the reproduction of colors by different displays and printers. (**B**) Stimuli plotted on a plane in perceptual color space (CIE LUV, u*= blue-yellow, v* = red-green) with gamut boundaries of the display shown by the white triangle.

**Table 1.**
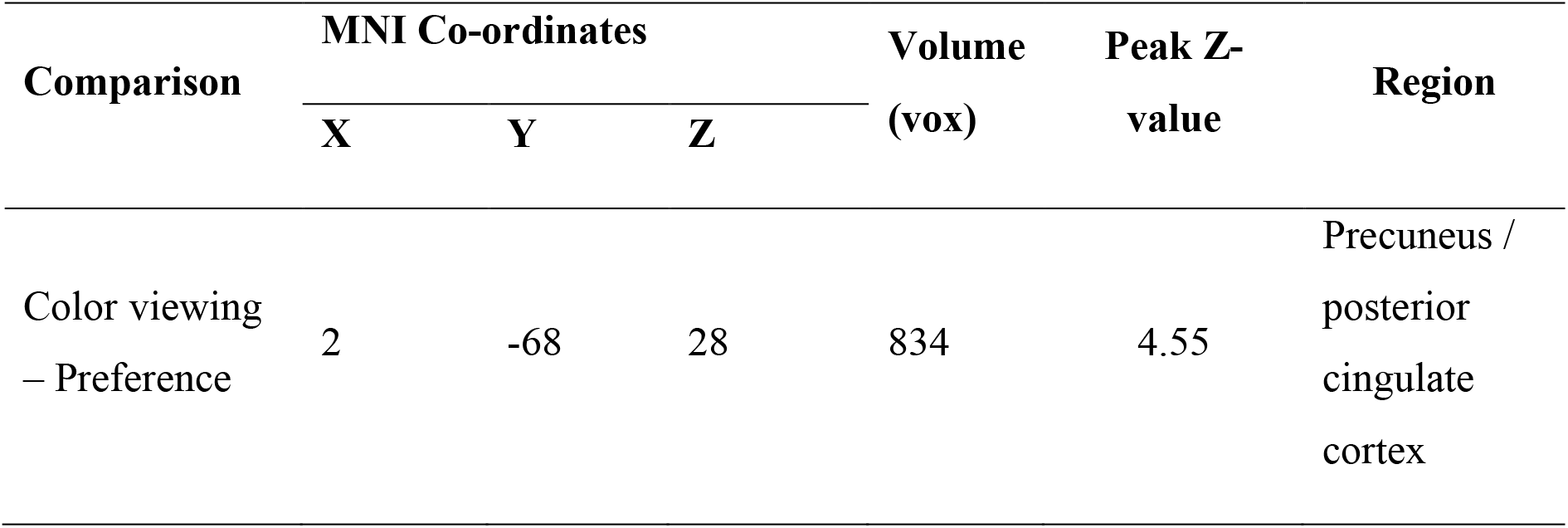
Preference related activation mean MNI coordinates (mm)

| Comparison | MNI Co-ordinates |  |  | Volume<br>(vox) | Peak Z-value | Region |
| --- | --- | --- | --- | --- | --- | --- |
|  | X | Y | Z |  |  |  |
| Color viewing<br>– Preference | 2 | -68 | 28 | 834 | 4.55 | Precuneus /<br>posterior<br>cingulate<br>cortex |

### Design and procedure

Participants completed 5 tasks whilst in the scanner: a task which required passive viewing of colors whilst judging the orientation of the stimuli (color viewing task); a color preference task where participants rated their preference of the stimuli; and three localizer tasks which aimed to localize color selective, object selective and color naming regions. The scanner was not active during the color preference task but this was conducted inside the scanner so that colors were rendered in an identical manner and seen in the same context as the passive color viewing fMRI task. Each task is outlined below.

### Color viewing scan

The aim of the color viewing task was to acquire BOLD response whilst participants viewed multiple presentations of the colored stimuli without making explicit judgments about the colors. Participants completed the task prior to any mention of color preference - during recruitment participants were told the broad aim of the study (to investigate the neural basis of color perception) and they were fully debriefed at the end of the study on the specific aim of understanding color preference. Participants were not required to make judgments about the colors, but were given an orientation judgement task in order to maintain attention on the stimuli. This involved judging whether the stimuli were one of four possible orientations (with a 1-4 button press, midline of the square stimulus angled −20°, −10°, 10° and 20° from vertical). Each colored stimulus was shown 7 times in a shuffled, pseudo-random order (203 trials), and the orientation of the stimuli was fully randomized. Stimuli appeared for 1000ms, and responses were recorded for a 1500ms window after stimulus onset. Stimuli were presented in a jittered-rapid event-related design. An optimal combination of stimulus order and Inter-trial interval (ITI) was generated with the make_random_timing.py script included with the AFNI package (Cox, 1996). We selected from 10,000 potential designs the ordering with the smallest amount of un-modeled variance. The task took 16 minutes and 20 seconds.

### Color preference task

As in Palmer and Schloss (2010), participants were first shown an array of all of the stimuli that subtended 9 degrees of visual angle and were asked to think about the color which they liked the most and the least. The aim of this was to familiarize participants to the range of the stimuli so that subsequent preference ratings could be made relative to the whole set. Next, each stimulus was presented individually and participants were asked to rate how much they liked the color on a scale from ‘not at all’ to ‘very much’ by sliding a cursor along a continuous response scale that was presented underneath the stimulus (as in Palmer & Schloss, 2010). The stimulus remained on the screen until participants made their response, with a 500ms intertrial interval. Stimuli were presented twice and in a pseudorandom order.

**Figure 2.**
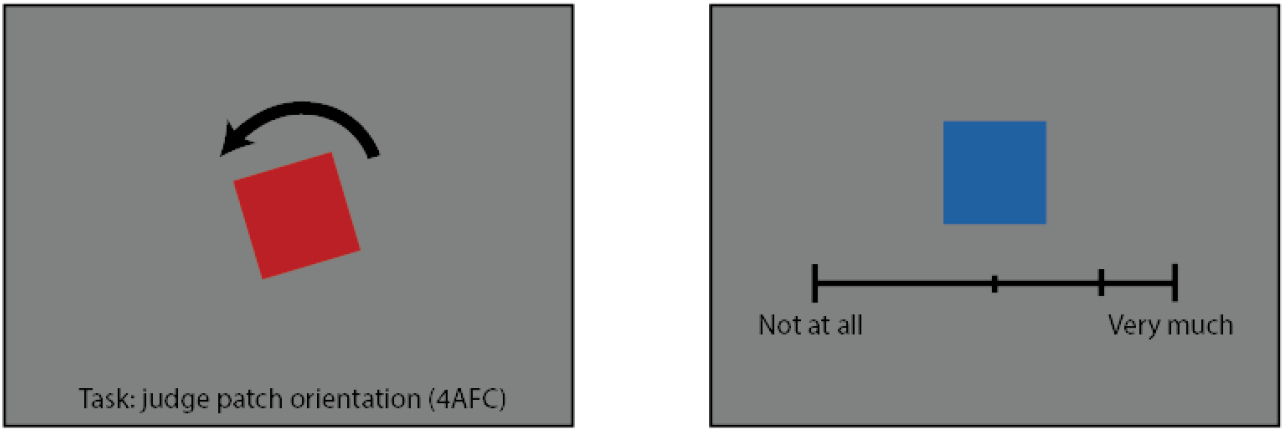
Experimental tasks. Left: Experimental task in color viewing scan, participants made an orientation judgment from 4 possible alternatives. Right: Example analogue color preference judgement in color preference task. In both cases, color patches subtended 2° of visual angle.

### Functional localizer scans

The three functional localizer scans (color selective, color naming selective and object selective) were block designs consisting of two conditions (ten blocks in total, five of each condition, duration = 4 minutes 30 seconds). Each localizer procedure is outlined below.

#### Color localizer

A standard procedure was used to localize color selective regions which contrasted BOLD response during viewing of gratings that were either a color and grey (color+grey) or were achromatic (light+dark) (Lafer-Sousa & Conway, 2013). Stimuli were full-field sinusoidal gratings (2.9 cycles per degree, drift rate 0.75 cycles per second, switching directions every 2s). There were five blocks of each type, lasting 10.48s each and separated by 10.48s fixation. Color+grey gratings were isoluminant for the standard observer (287.1 cd/m^2^) and consisted of D65 grey and had the chromaticity of the Red and Purple from the Saturated set and the Orange, Yellow and Green stimulus from the Light set. Light+dark gratings were made up of d65 grey at two lightness levels (L*=91.1, and 151.3). Participants were instructed to pay attention to a black fixation cross that was centrally presented throughout the task and to respond when one of the arms of the fixation cross disappeared (1-2 targets per block).

#### Object localizer

A standard object localizer procedure was used to localize the object selective Lateral Occipital Complex ((LOC; Grill-Spector, Kourtzi, & Kanwisher, 2001). The two conditions of the object localizer task were the presentation of either intact- or scrambled-objects (presented in black and white and compiled from several online object image repositories and prepared with Matlab, The Mathworks Inc., 2012). Each image was displayed for 2000ms each and separated by 500ms of fixation cross, and each block contained 10 images. Participants were instructed to pay attention to the images and to respond whenever a red dot appeared on any image (1-2 targets per block).

#### Naming localizer

The naming localizer contrasts internally voiced color naming with internal irrelevant voicing in order to localize brain regions associated with active color naming. The two conditions differed only in the task instructions, in one condition, participants were instructed to use internal voicing to freely name the color of each stimulus whilst the color was shown, in the other condition they were instructed to silently repeat the irrelevant obscure color term ‘Tan’. Participants were informed of the condition type by the fixation cross changing to either a ‘C’ or a ‘T’ for 3700ms prior to the onset of a block. Each stimulus was shown for 1300ms, with 200ms of fixation cross and condition reminder during the inter-trial interval. Stimuli were pseudorandomly chosen from the full stimulus set and there were eight stimuli per block.

### Data acquisition

All images were acquired on a 1.5 T Siemens Avanto scanner using a 32-channel phased-array head coil. Participants were placed in the scanner in a supine position. Functional images were acquired using a T2*-weighted gradient-echo EPI sequence (TR = 2620 ms, TE = 42, flip angle = 90, FOV = 192 × 192 mm, matrix = 64 × 64). Each functional volume consisted of 35 contiguous 3.6 mm thick axial slices with 3 × 3 mm in-plane resolution. In addition, a high resolution (1 mm3) T1-weighted whole brain anatomical volume was collected with a magnetization-prepared rapid gradient-echo (MPRAGE) sequence for purposes of co-registration and standardization to a template brain. Finally, a field map was collected to allow for correction of geometric distortions induced by field inhomogeneities.

### Data pre-processing and analysis

Data pre-processing and analysis were performed using the Oxford Centre for Functional Magnetic Resonance Imaging of the Brain (FMRIB) Software Library (FSL 5.0.8, www.fmrib.ox.ac.uk/fsl). Pre-processing was performed using FEAT (fMRI Expert Analysis Tool, version 6.00). The initial four volumes of data from each scan were removed to minimize the effects of magnetic saturation. Motion correction was followed by spatial smoothing (Gaussian, FWHM 8mm) and temporal high-pass filtering (cutoff, 0.01 Hz). B0 unwarping was performed using FUGUE (http://www.fmrib.ox.ac.uk/fsl/fugue/index.html). For each EPI run, non-brain data were removed using the FMRIB Brain Extraction Tool (Jenkinson, Bannister, Brady, & Smith, 2002). Registration of the functional data followed a 2-stage process using linear registration with the same FMRIB tool: each functional run was first registered to the high resolution T1-weighted MPRAGE image (7 degrees of freedom), and then registered to the Montreal Neurological Institute (MNI) 152 standard template anatomical image (12 degrees of freedom).

The BOLD signal was modelled by convolving the predictor function of event timing with a standard model of the hemodynamic response function (HRF) for each of the 29 color conditions (regressors). For each individual participant, a unique contrast vector was computed from the preference and naming RT behavioral measures. Ratings and scores were converted to z-scores relative to the individuals’ mean value ([x^i^-mean(x)]/stdev(x)), and then normalized by dividing by the max value such that each vector has a mean of 0. These contrasts reflect the extent to which the BOLD response in each voxel correlates with subjective color preference and individual color naming response time. In addition to the above subjective contrast vectors three general contrasts were included, a contrast of all events regardless of color compared to fixation baseline (active > rest), and two colorimetric repressors reflecting the normalized chroma and lightness of each color condition. Group level analysis was carried out using the FMRIB Local Analysis of Mixed Effects tool (Beckmann, Jenkinson, & Smith, 2003). Resulting Z statistic images were thresholded using a cluster-forming threshold of Z > 3.1, and a familywise error (FWE) corrected cluster extent threshold of p < 0.05, based on the theory of Gaussian Random Fields. Cortical labels in the whole brain analyses were determined using the Harvard-Oxford Cortical Structural Probabilistic Atlas (Desikan et al., 2006).

Functional localizer regions of interest (ROIs) were individually defined in each participant using each of the localizer scan contrasts (chromatic > achromatic gratings; intact > scrambled objects; internal color naming > internal irrelevant voicing). At the first level, the resulting statistical maps were thresholded at a minimum threshold of p > 0.001 (uncorrected) for each of the three localizer scan types. Localizer contrasts were also taken to the group level using a cluster-forming threshold of Z > 3.1, and a familywise error (FWE) corrected cluster extent threshold of p < 0.05 in order to generate group level masks for each localizer. Group level masks were transformed to each participants’ native brain space and used to further constrain individual level masks. All subjects showed sufficient activation to generate localizer masks for each of the three scan types. For each region mask in each individual, the contrast of parameter estimates (COPE) for all contrasts of interest were extracted using the FMRIB Featquery tool and converted to units of percent signal change. The extracted values are measures of effect size, and in the instance of subjective contrast vectors of preference and naming RT, reflect the extent to which a given region correlates with that behavioral measure. In order to determine group level effects the extracted region effect size measures were averaged across participants and each region subjected to a one-sample t-test against a mean of zero.

## Results

### Color preference curves

Figure 3 shows color preference plotted as a function of hue. The most preferred colors include saturated blues and greens and the least preferred include dark yellow and dark orange.

**Figure 3.**
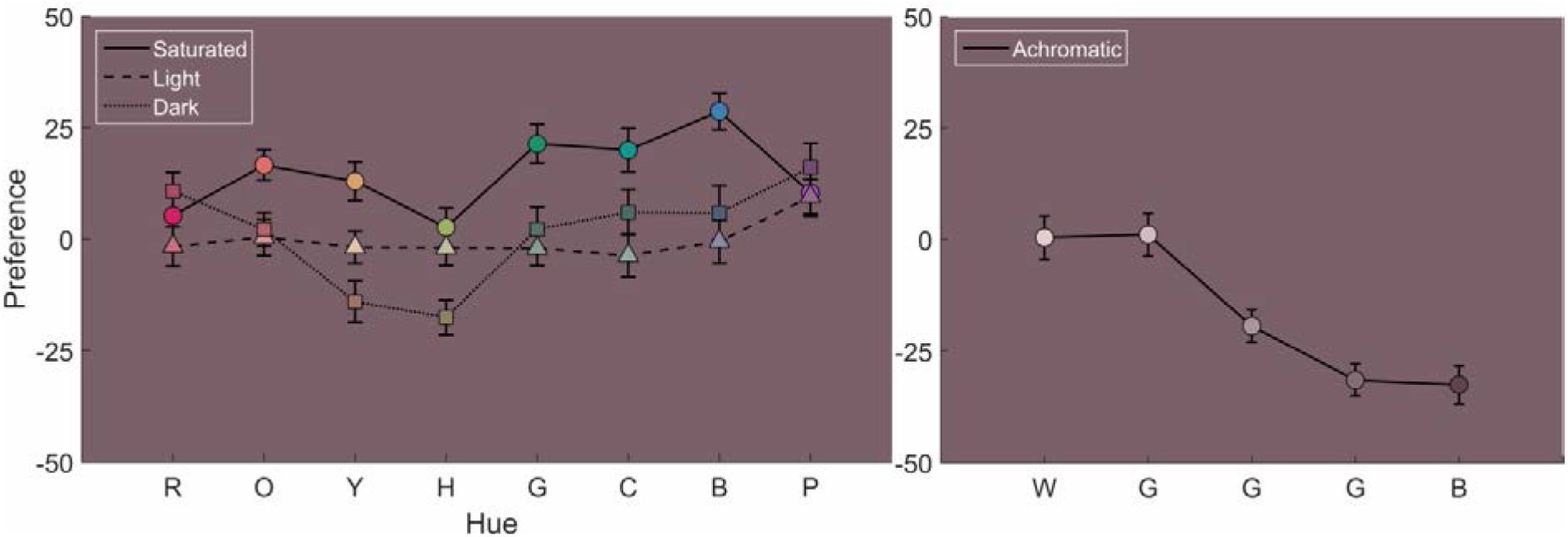
Mean preference ratings (±SEM) for light, saturated and dark stimuli (left), and achromatic stimuli (right). The X-axis gives the hue; red (R), orange (O), yellow (Y), chartreuse (H), green (G), cyan (C), blue (B), purple (P), white (W), grey 1-3 (G), black (B). The Y-axis shows arbitrary values assigned to analogue preference ratings from color preference task which used a scale where the endpoints were marked ‘not at all’ (−50) or ‘very much’ (+50). The marker colors are for illustrative purposes only and are an approximation of those in the experiment.

Individual measures of color preference were reliable and consistent: preference ratings showed significant correlations between the first and second time each color was rated for each individual (average r = 0.78). All individual correlations were positive and significant at p <0.005.

### Whole brain analysis of color preference

In order to determine the brain regions which correlate with color preference we conducted analyses to determine which regions of the brain have a BOLD response when viewing colors that correlates with participants’ color preference ratings.

First, we checked that participants were paying attention during the color viewing task. The mean performance at judging the orientation of the patches in the color viewing task was 86% correct (sd = 11.5%) which is comfortably above chance levels (25%). One participant was excluded from all analyses due to poor performance on this task, performing below two standard deviations from the mean accuracy. This participant also performed at chance levels on other behavioral tasks in the study.

Second, a whole brain correlation was computed between the mean BOLD response for each of the 29 colors during the color viewing task and each individual’s color preference ratings. Figure 4 shows clusters of voxels (blue) with a significant positive correlation with color preference. The peak of the activation is in the precuneus cortex (Peak MNI coordinates: 2, −70, 30, size: 487 voxels, peak Z: 4.7). The activation spans across the posterior midline cortex into both hemispheres and extends from the precuneus cortex into the posterior cingulate gyrus.

**Figure 4.**
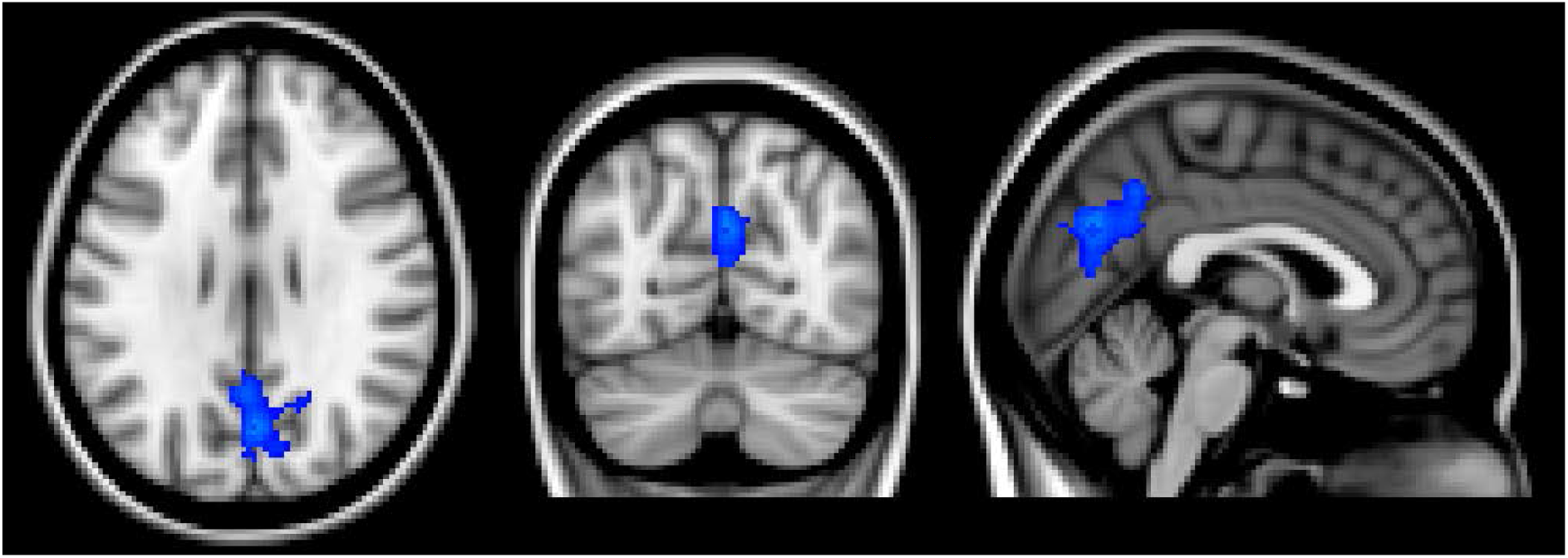
Whole brain color preference correlation with BOLD when colors are viewed. Images are thresholded with a cluster-forming threshold of Z > 3.1, and a familywise error (FWE) corrected cluster extent threshold of p < 0.05. Peak MNI coordinates: 2, −70, 30, size: 487 voxels, peak Z: 4.7. Images are shown in neurological convention (left hemisphere on the left).

### Functionally defined Regions of Interest

In order to investigate previously established relationships of color preference with measures such as color naming, color-object associations and sensory encoding of color, three short functional localizer scans for color selectivity, object selectivity and color naming were analyzed. Simple contrasts in each of these scans (chromatic > achromatic gratings; intact > scrambled objects; internal color naming > internal irrelevant voicing) were used to define ROI masks for each individual subject (see methods). Figure 5 shows the group level localizer masks for all regions in MNI space. The peak MNI coordinates for all ROI’s are shown in table 2. Bilateral object selective regions from the object localizer were found in the inferior lateral occipital cortices in all participants, these regions were labelled as left and right LOC. In the color selectivity localizer, bilateral regions were found within the occipital fusiform gyrus in all participants and labelled as left and right Fus. The color naming localizer produced two distinct left lateralized frontal regions, the peak of the medial region was in the paracingulate gyrus, this region was labelled ParaC. The peak of the lateral region was in the precentral gyrus and labelled PreCG. Both naming localizer masks were located in all participants except one, who failed to show sufficient activation to define a ParaC mask.

**Figure 5.**
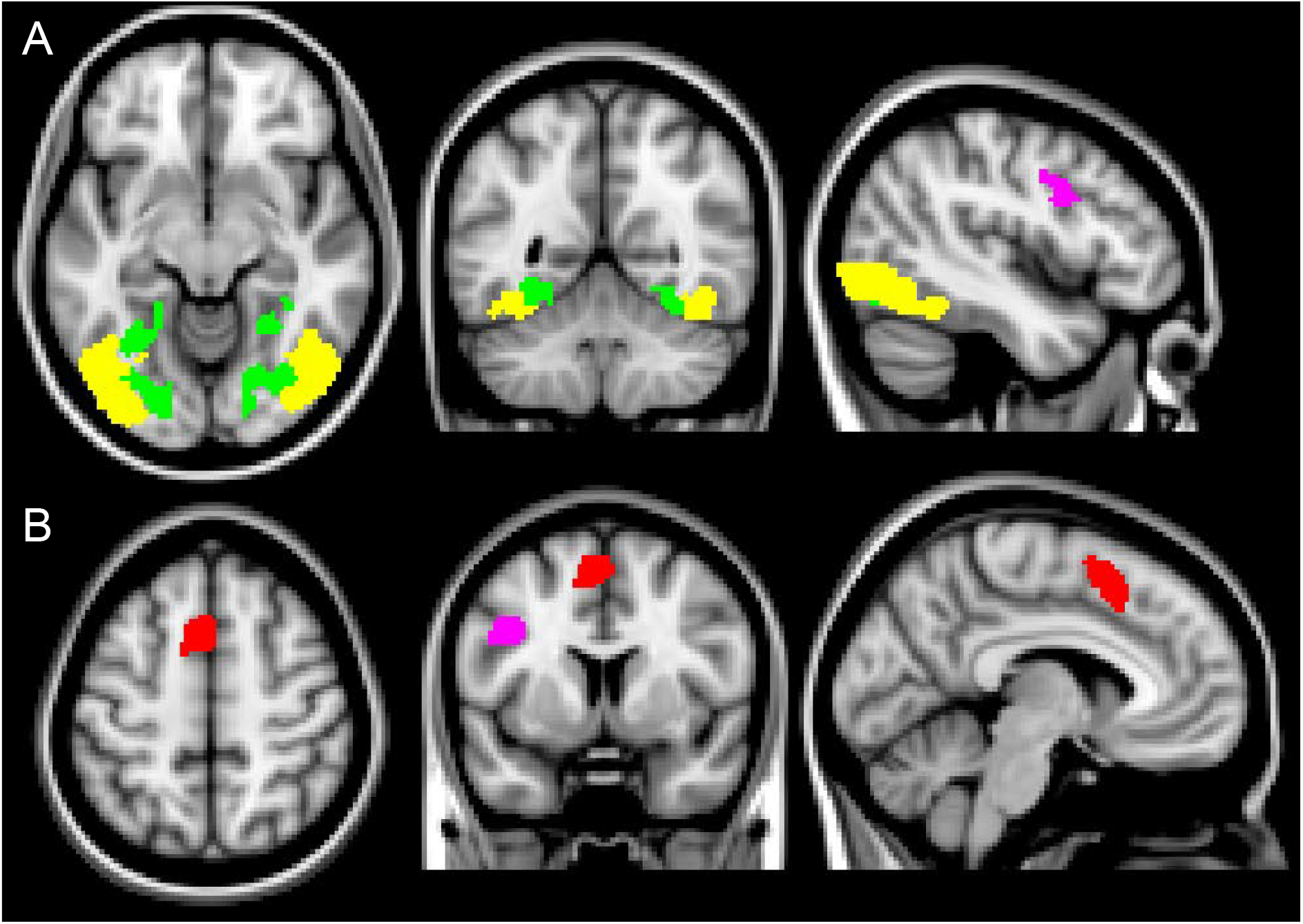
Location of all group level localizer masks displayed in MNI space. (A) shows object selective regions (Yellow, LOC) and color selective regions (Green, Fus). (B) shows naming localizer regions (Red, ParaC; Pink, PreCG). ROIs were defined at the individual level from independent functional localizer scans. Images are shown in neurological convention (left hemisphere on the left).

**Table 2.** Mean MNI coordinates (mm) of the center of gravity of all localizer masks across individuals. All ROIs were defined in individual EPI space and transformed into MNI. Standard error is reported in parentheses.

| **Region** | **N** | **X** | **Y** | **Z** | **Volume(vox)** |
| --- | --- | --- | --- | --- | --- |
| Fus (Color selective) |  |  |  |  |  |
| L | 20 | -28 (0.05) | -68 (0.07) | -14 (0.04) | 690 (15.66) |
| R | 20 | 27 (0.06) | -69 (0.1) | -12 (0.03) | 663 (14.58) |
| LOC (Object selective) |  |  |  |  |  |
| L | 20 | -38 (0.03) | -69 (0.06) | -11 (0.05) | 935 (18.28) |
| R | 20 | 42 (0.04) | -67 (0.04) | -10 (0.03) | 739 (15.66) |
| Color naming |  |  |  |  |  |
| PreCG | 19 | -42 (0.07) | 5 (0.06) | 32 (0.07) | 270 (5.6) |
| ParaC | 20 | -3 (0.06) | 14 (0.06) | 51 (0.05) | 331 (7.33) |

In the object localizer and color selectivity localizer participants’ undertook irrelevant tasks designed to maintain alertness and attention on the stimuli and mean performance in these tasks was 92% correct (sd = 7.9%) and 70% correct (sd = 10.8%) respectively. All participants performed well above chance in these tasks.

There was no difference in the pattern of response between the right and left hemispheres for the object selective and color selective, bilateral ROIs. Accordingly, all subsequent analyses were based on a pooled analysis in which ROIs from the right and left hemispheres were combined. The naming localizer produced two functionally distinct regions which were analyzed separately.

For each region mask in each participant’s individual EPI space, the parameter estimates were extracted for correlations with preference. The extracted parameter estimates are measures of effect size which reflect the strength of the relationship between preference and BOLD (See figure 6). One-sample t-tests were conducted on the parameter estimates and identified that estimates in Fus, PreCG and ParaC regions were all not significantly different to 0 (largest t=.35, smallest p=.73), but the LOC activation reached significance with a one-tailed test (t(19)=1.85, p=.084). Bayes Factors calculated with a cauchy prior of 0.707 (Wetzels & Wagenmakers, 2012; Wagenmakers et al, 2018) identified support for the null for the Fus (B = 0.246), PreCG (B= 0.256), & ParaC (B= 0.232) regions, and did not provide support for either the null or alternative hypothesis for the LOC (B: 0.863).

**Figure 6.**
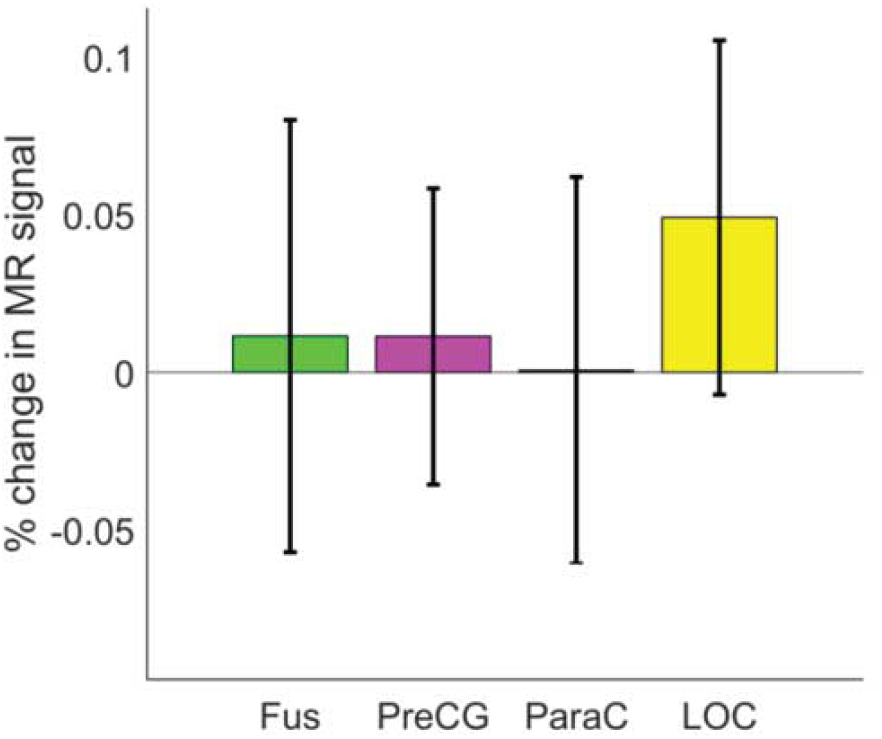
Preference correlations in all ROIs during passive color viewing. Bars denote functionally defined ROIs, Green=color selective Fus, pink and red=naming responsive ParaC and PreCG respectively, yellow=object selective LOC. See figure 5 for anatomical depictions of these regions. Error bars represent 95% confidence intervals.

### Colorimetric contributions to color preference

Perceptual dimensions of color such as chroma are known to be related to color preference (Guilford & Smith, 1959; Palmer & Schloss, 2010). Therefore, correlations of chroma and lightness (CIE LCH) with color preference and BOLD were computed in order to assess the extent to which colorimetric variation in the stimulus might account for color preference. A significant positive relationship was not found between color preference and lightness (r= 0.14, p=0.47), but was with CIE chroma (r = 0.42; p < 0.05). See figure 7 for scatter plots of these relationships.

**Figure 7.**
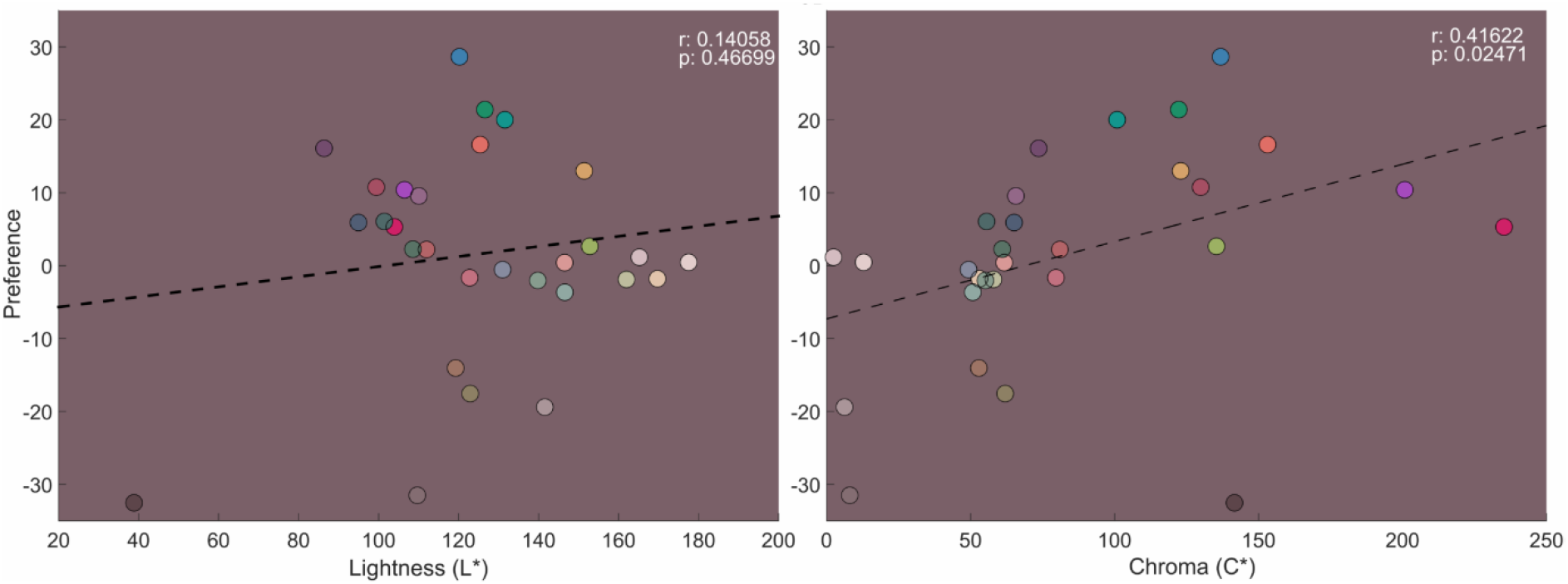
Correlation between colorimetric measures CIE lightness (L*) and CIE chroma (C*) and preference for each color. Preference measure is averaged by color across participants.

Whole brain colorimetric regressors of both lightness (L*) and chroma (C*) were included in analyses of the color viewing scan in order to look for voxels which show a significant correlation with these perceptual properties of color. A region with a significant positive correlation with lightness was found (see figure 8). The peak of the activation is in the cuneal cortex (Peak MNI coordinates: −12, −70, 18, size: 1003 voxels, peak Z: 5.33). The activation spans across the midline into both hemispheres, shows minimal overlap with the precuneus preference related activations and is confined mostly to the occipital lobe.

**Figure 8.**
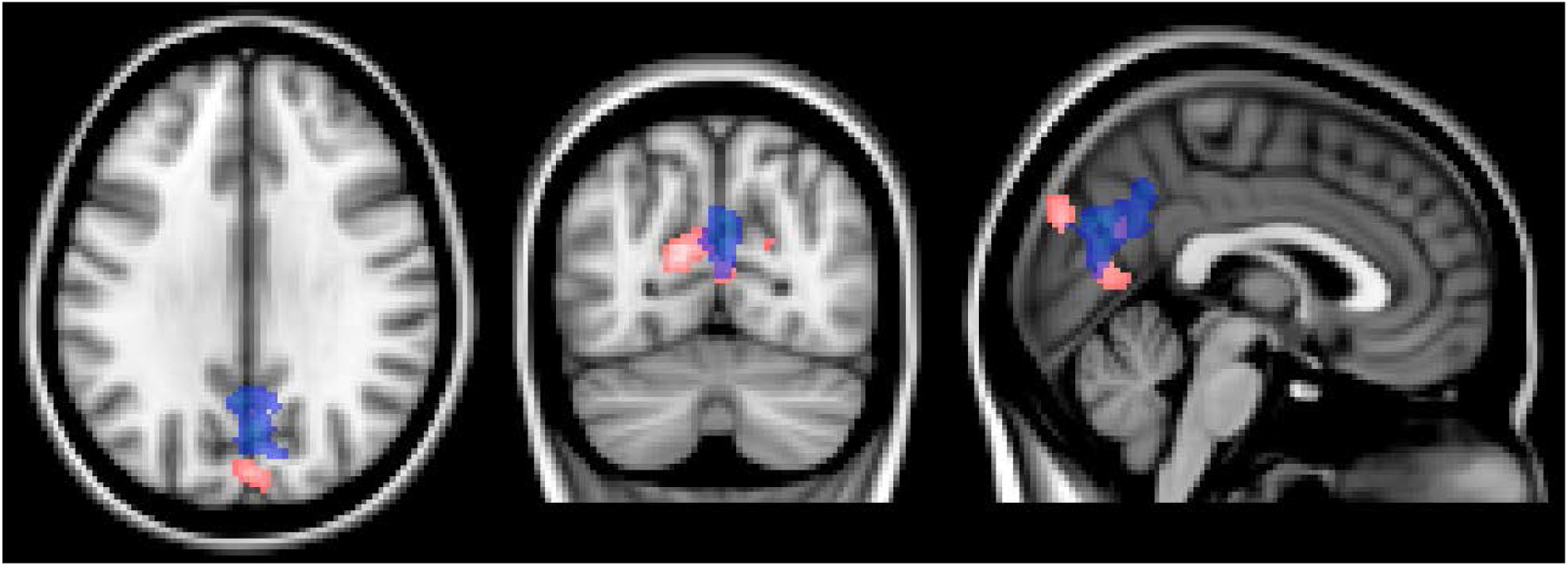
Whole brain correlation of CIE lightness (L*) with BOLD from the color viewing task (pink). Translucent blue shows position of precuneus preference correlation from figure X (color viewing scan), Images are thresholded with a cluster-forming threshold of Z > 3.1, and a familywise error (FWE) corrected cluster extent threshold of p < 0.05. Peak MNI coordinates of lightness correlation: −12, −70, 18, size: 1003 voxels, peak Z: 5.33. Images are shown in neurological convention (left hemisphere on the left).

## Discussion

The current study aimed to identify brain regions associated with color preference, by correlating BOLD activity when people passively view colors with their color preferences measured after the scan. We had two sets of predictions: one set which was based on previously established behavioural relationships of color preference with color naming, object perception and sensory encoding of color (e.g., Àlvaro et al., 2015; Hurlbert & Ling, 2007; Palmer & Schloss, 2010), and another set based on prior neuroimaging studies of value and aesthetics for stimuli other than color (e.g., Grueschow et al., 2015; Vessel et al., 2013). We localised color naming, object selective and color selective regions of cortex in order to test for correlations of BOLD activity and preference in the regions that relate to our first set of predictions, and we also conducted a whole brain analysis in order to investigate the network of regions identified in prior neuroimaging studies of value and aesthetics. Color preference did not correlate with activity in localised color naming regions or color selective regions of visual cortex, but there was a marginally significant effect of color preference in a localised object processing region LOC. The whole brain analysis found a significant correlation between color preference and activity in the posterior midline cortex (PMC), centred on the precuneus but extending into the adjacent posterior cingulate and cuneus. This was outside of the regions of interest identified with the three functional localiser tasks. The PMC effect could not be accounted for by lightness or chroma since only lightness was correlated with activity and this effect was almost entirely in the occipital lobe.

First, we discuss the main result of a correlation in the PMC between BOLD activity when colors are passively viewed and how much people like those colors. Prior studies have revealed a role of the PMC in value based processing. For example, a meta-analysis of the brain regions that compute subjective value identified several regions, including a ventral PCC effect that overlaps with the PMC region identified in our study (Clithero & Rangel, 2014). A specific role for this region in value processing was identified by Grueschow et al., (2015) who showed that preference ratings for movies correlated with activity in the PMC while the covers of the movie DVDs were presented. Importantly, this correlation was present on trials when the subjective value of the movie was irrelevant to the ongoing task (which was to count the number of faces in the picture). The authors suggested that, “The automatic nature of SV [“subjective value”] representations in PCC may constitute an important evolutionary advantage, as it could ensure that SVs of external environmental features are continuously encoded with minimal use of attentional resources”. Prior studies have revealed a role of the PCC in automatic encoding of subjective value for stimuli that are obviously value-laden such as movies (Grueschow et al., 2015), subjective value for monetary reward (Kable & Glimcher, 2007), or paintings (Jacobsen, Schubotz, Höfel, & Cramon, 2006; Vessel, Starr, & Rubin, 2012). Unlike monetary reward, art or higher level configural stimuli (e.g. movie posters), we consider abstract color patches less likely to elicit conscious value judgments in the absence of instruction to evaluate them. Here we show that the PCC is automatically encoding subjective value even for basic features of the environment such as color.

The PMC is most strongly associated with being the main “hub” of the DMN, a network of regions that are activated while participants are engaged in internally generated thought compared with external attention (e.g. while mind-wandering (Buckner, Andrews-Hanna, & Schacter, 2008; Mason et al., 2007), self-referential processing (Northoff et al., 2006), autobiographical memory retrieval (Philippi, Tranel, Duff, & Rudrauf, 2015), and thinking about the future (Kable & Glimcher, 2007; Levy & Glimcher, 2011). One prior study which found that DMN activation was greater the more that images of art were liked, suggested that this was due to the DMN’s role in self-referential processing and that it provided evidence that liked stimuli resonate more with a sense of self and self-identity than disliked stimuli (Vessel et al., 2013). The involvement of the DMN in color preference that has been identified in the current study could potentially therefore support the idea that self-identity shapes color preferences, an idea that is consistent with evidence that sex stereotyped color preferences emerge around the time that gender identity develops (LoBue & deLoache, 2011).

The PMC is also strongly implicated in semantic processing (e.g. Binder et al., 2009) and has been identified as supporting amodal representations of objects (Fairhall & Caramazza, 2009). For example, in one study, patterns of activity in the PMC were similar when an object was presented in either word or picture form, suggesting that it represented the core “concept” of the object as opposed to the specific stimulus present (Fairhall & Caramazza, 2009). By contrast, regions that showed similarity for just the picture presentations of objects overlapped our LOC region of interest. Therefore, whilst we do not find an effect in the LOC, we do find an effect in a region involved in higher order object processing. This correlation between color preference and activation in the PMC could provide support for the theory that color preferences are related to color-object associations (e.g., Palmer & Schloss, 2010). Further research which localises the brain regions specifically responsible for color-object associations, and which identifies whether these regions overlap with our PMC effect would provide further support for this position.

Whilst we find a strong correlation of preference and activation in the PMC, we do not find correlations in other regions associated with value and aesthetics in prior neuroimaging studies of other stimuli such as the orbitofrontal cortex, or regions predicted by other theories of color preference such as color naming or color selective regions. The lack of effects in the orbitofrontal cortex could well support the argument that this region is important when observers make explicit judgments of value or aesthetics rather than the automatic preferences measured here (see Conway & Rehding, 2013). The lack of effects in color naming or color selective regions should not be taken as evidence against the theories that color preference is related to color naming (e.g., Álvaro, Moreira, Lillo, & Franklin, 2015) or the sensory mechanisms (Hurlbert & Ling, 2007) of color preference. These relationships have been established in prior behavioural studies, and their neural basis may lie in other regions of the brain that were not captured by our functional localisers.

For many decades, scientists and artists have wondered why simple patches of color have valence and ‘hold’ affect (Zajonc, 1980). Here we present the first neuroimaging study which provides leverage in addressing this question. We found that PMC activation is greater the more a viewed color is liked even in the absence of an explicit preference judgment or context. This suggests that color preferences are registered by the brain automatically, suggesting that color preference may be a pervasive aspect of the visual processing of scenes. The findings have implications for understanding the neural basis of value or aesthetics beyond color, by identifying that the PMC is related to automatic encoding of subjective value even for basic visual features. Future studies may establish why the human brain automatically computes value for basic visual features and identify the perceptual, cognitive and behavioural consequences of these automatic preferences.

## Acknowledgements

This research was supported by a European Research Council grant to A.F. (project CATEGORIES: ref 283605).

